# Noise in Competing Representations Determines the Direction of Memory Biases

**DOI:** 10.64898/2025.12.17.694673

**Authors:** Andrey Chetverikov, Sabrina Hansmann-Roth

**Author notes:** Correspondence should be addressed to: Department of Psychosocial Science, Faculty of Psychology, University of Bergen, Post box 7807, 5020 Bergen, Norway.

## Abstract

Memories are reconstructions, prone to errors. Historically considered a nuisance, memory errors have recently gained attention when found to be systematically shifted away from or toward non-reported items, promising insights into memory mechanisms. We propose that these biases are optimal and inevitable when the brain disentangles overlapping memory signals, predicting that bias direction depends on the noise distribution between memorized items, not just absolute noise levels. We tested this prediction in four color-memory experiments using novel stimuli with independently varied noise levels. The results support our hypothesis: targets with the same absolute noise level can be repelled from or attracted to non-target items, depending on their relative noise levels. We further show that the model can fit nonlinear bias patterns observed in human data with noise levels as the only free parameters. These findings challenge currently dominant models and support signal disentanglement as a unifying explanation of memory biases.

---

How do we remember visual information? For a long time, the visual short-term memory literature focused on the capacity and fidelity of memory, leading to the common assumption that individual items in memory are isolated from each other. For example, a widely used mixture model of working memory assumes that the reported item representation is not affected by non-reported items, with the exception of possible catastrophic failures leading to swap errors ^1^. However, no memory is an island. The last decade has brought an abundance of studies demonstrating interactions between memory representations and between memory and perception ^2–13^. These interactions result in systematic shifts – or biases – in behavioral responses when analyzed as a function of non-target item features. Similarly, neuroimaging studies demonstrate shifts in decoded representations of memorized items under the influence of other items ^14–18^. While such biases are by now well documented, their theoretical rationale remains elusive.

A currently dominant explanation suggests that attractive biases, where an item’s report shifts towards non-targets, arise from confusion or purposeful integration of information from multiple items ^5,7,19,20^. This is more likely when representations are noisy ^5,13,21^, and indeed, Lively et al. ^9^ demonstrated that increasing working memory load leads to stronger attractive biases. Repulsive biases, where an item’s report shifts away from non-targets, are further explained ^4,5,21^ by the brain’s attempt to reduce interference between representations by shifting them away from each other, a phenomenon that is especially pronounced when items are similar and hence more confusable. According to this perspective, the *absolute* amount of noise in target and non-target representations governs the bias direction: similar but distinguishable items are repelled, while poorly distinguishable ones are attracted. Furthermore, these theories assume that biases have the same direction for affected items: either both are attracted due to confusion or integration, or both are repelled from each other to reduce interference.

We recently proposed an alternative explanation of memory biases (Figure 1A) based on the observation that the brain must disentangle signals from multiple stimuli to infer their properties^22^. Notably, this normative model explains both repulsive and attractive biases with a single mechanism, showing that they emerge as optimal solutions to the problem of mixed signals under varying noise and stimulus similarity. That is, not only is the amount of noise in memory important (for example, under varying memory load), but also how noise is distributed between the items, specifically how much noisier the target item is compared to a competing non-target. The model further predicts that, under unequal noise, the biases for the affected items can have opposing directions, again contradicting currently dominant theories.

**Figure 1.**
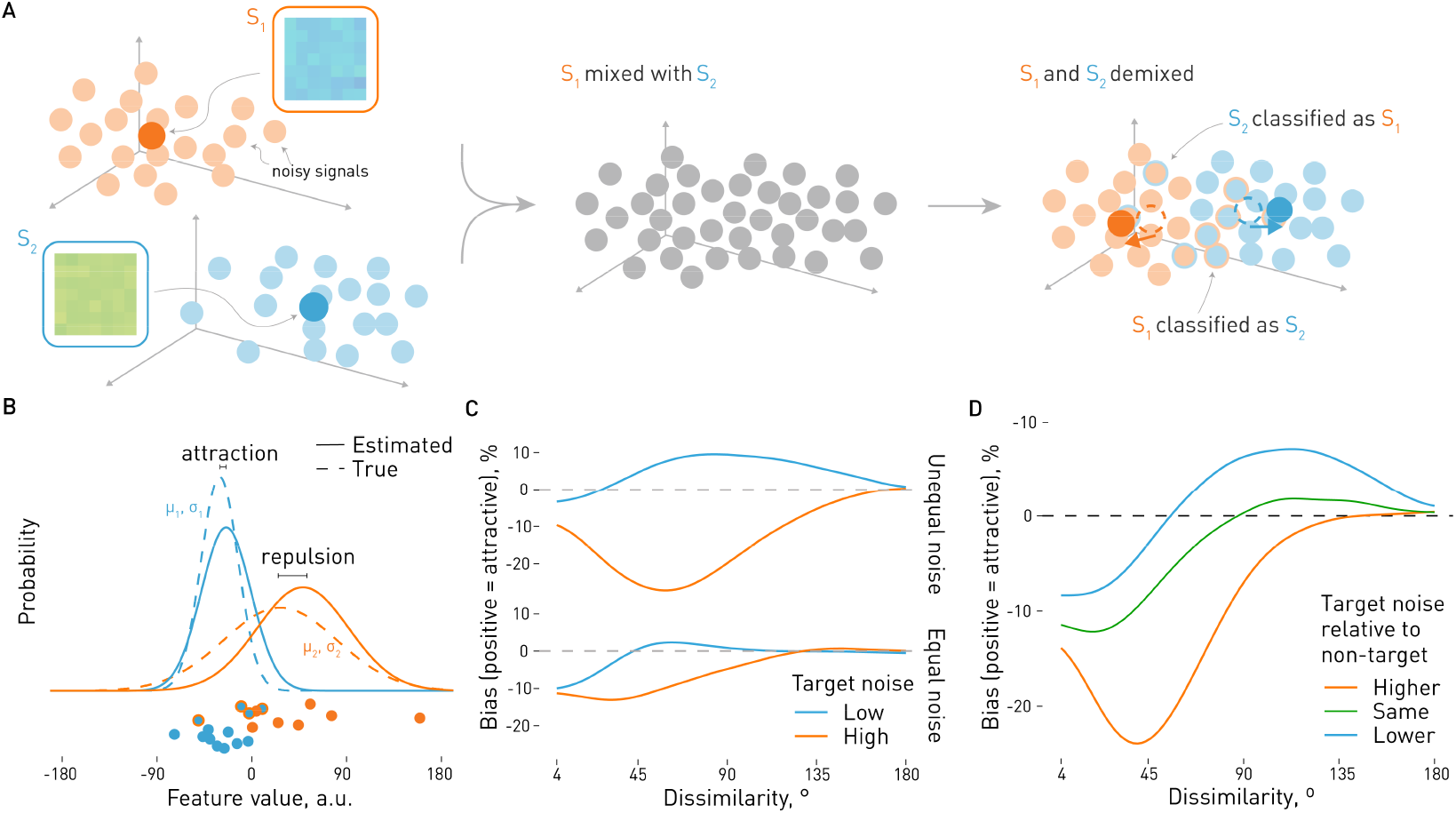
Predictions of the Demixing Model. **A**. Model outline. The Demixing Model assumes that the observer receives noisy signals from multiple items (color patches S1 and S2, left). These signals are mixed in a common representational space (middle), so stimulus features (e.g., color) must be reconstructed by attributing signals to items, i.e., “demixing” them. Even an optimal solution is imperfect: some signals generated by one item are attributed to the other (right; dot outline – true source, fill color – attributed source; uncertainty in assignments is omitted for clarity). As a result, the estimated stimulus values (dashed outlines) systematically shift relative to the true ones (opaque circles), producing behavioral biases. **B-D**. Non-trivial predictions of the model emerge when the two stimuli differ in noise: the less noisy item is attracted toward the noisier one, while the noisier item is repelled away from the less noisy one. B. These predictions can be intuitively understood by considering the overlapping distributions of signals coming from the two stimuli (dashed lines). Samples (dots) from the noisier stimulus (orange) that fall close to samples from the less noisy one (blue) are likely to be attributed to the less noisy item (fill color – attributed source, outline – true source). As a result, the estimated distributions (solid lines) shift: the noisier stimulus is repelled from the less noisy one, while the latter is attracted to the former. **C**. The resulting average bias pattern across different similarity levels shows more attraction for the less noisy stimulus and more repulsion for the noisier one when the noise levels of the two items are unequal (“unequal noise”). This difference is more pronounced than when the target and non-target have the same noise level (“equal noise”). **D**. When the target noise level is kept constant, varying the non-target noise level leads to more attraction or repulsion depending on whether the target has a lower or a higher noise level relative to the non-target. Note that the absolute magnitude and direction of biases also depend on other factors (see Methods and Fig. S1).

Here, we test this model using a novel visual working memory task with color stimuli varying in noise. We first show, in simulations, specific predictions for: (1) a case where the noise of the target and non-target items is either swapped or simultaneously changed for both items; and (2) a case where only non-target noise changes. We then demonstrate in four experiments that the relative amount of noise in the remembered stimuli strongly affects the magnitude and direction of memory biases, contrary to the common view but in line with the model’s predictions. Finally, we use neural networks to interpolate the simulated model predictions and fit the model to human data, demonstrating that, despite having just a few parameters, the model can reproduce complex nonlinear patterns of biases as a function of noise and similarity between the stimuli.

## Results

### Simulated model predictions

We first simulated the predictions of the Demixing Model^22^ for color reports with two stimuli. The model represents the stimuli as points in a two-dimensional space where one axis corresponds to the reported feature and the other to a projection of all other features to the extent that they allow separation of item identities. For example, in memory studies, the stimuli are commonly presented at different locations or timepoints, so the second dimension of the model would correspond to space or time, respectively. Each stimulus is characterized by a noise distribution that we modeled as the product of a wrapped normal distribution (corresponding to the circular color dimension) and a normal distribution (corresponding to the identifiability dimension). The observer obtains multiple samples (estimates of each stimulus’ features) from both stimuli with equal probability:

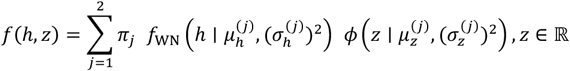

where *j* ∈ {1,2} indexes the two mixture components, each corresponding to one of the stimuli, *h* ∈ [0,360) is the sample’s hue with the mean of 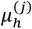 and the standard deviation of 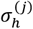 for each stimulus, *z* is the identifiability dimension of the samples with mean 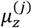 and the standard deviation of 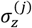, and *σ*_*j*_ are the mixture weights (equal to 0.5 for both stimuli), *f*_WN_ is the wrapped normal and *ϕ* is the normal distribution function.

The predictions of the model were obtained for different noise levels of the stimuli and different similarity levels between the stimuli in the color domain through simulations (see Methods for details). For each combination of the noise parameters, we simulated *N* = 20 observations and found the best estimate of the parameters 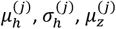 and 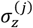 by fitting a Gaussian mixture model to the simulated data. This procedure was repeated 10000 times for each 4° step of the similarity between the stimuli in the color domain to estimate the probability of different biases (i.e., obtain a likelihood surface of the model predictions). Then, the asymmetry in probabilities of positive (towards the other item) and negative (away from the other item) errors was computed for each similarity step to match the metric used in the experiments.

We confirmed the earlier observation^22^ of a non-trivial bias pattern when the two stimuli had unequal noise (Figure 1B-D). In particular, the less noisy item can be attracted to the noisier one at the same time that the noisier item is repelled from the less noisy one (Figure 1C). This asymmetry can be intuitively understood by considering that the samples from the noisier item can by chance be similar to those from the less noisy item, and are thus attributed to the same source (Figure 1B). Because the noisier item has ‘lost’ these samples, the estimate of its value (i.e., color) will shift away from the less noisy item, while the estimate of the less noisy item will shift towards the noisier one.

The diverging bias pattern cannot be explained solely by the difference in the target noise level, as shown by comparison with the ‘equal noise’ case in Figure 1C. With equal noise, the target and non-target (i.e., the non-reported item) have the same noise levels, and the difference due to a change in target noise is much weaker. Furthermore, additional simulations showed that the overall bias pattern might shift towards more attraction or more repulsion, but we still expect the interaction between target noise and noise equality (Figure S1).

We also found that, due to the same asymmetry, it is possible to shift the bias, potentially changing its sign, by changing the non-target noise level (Figure 1D). That is, a target can be attracted more to (repelled less from) the non-target when the target noise is lower than the non-target noise, compared to the same target in the presence of a less noisy non-target. This pattern has the same underlying logic as the one described above (Figure 1C; when target has higher noise than the non-target, it means that it is the noisier item, and vice versa), but demonstrates that changes in the target noise are not necessarily needed to change the bias sign.

We then tested the model predictions: Experiments 1-3 changed both target and non-target noise levels focusing on predictions in Figure 1C, while Experiment 4 tested predictions in Figure 1D.

### Experiments 1 & 2

In Experiments 1 and 2, observers performed a visual working memory task with novel mosaic stimuli consisting of numerous small colored squares that together defined the overall stimulus hue (Figure 2A). Such stimuli are commonly used in ensemble perception studies showing that observers can effortlessly extract their properties, such as the average hue or hue variability^23,24^. At the same time, it allows precise manipulation of the amount of noise in memory for an individual stimulus.

**Figure 2.**
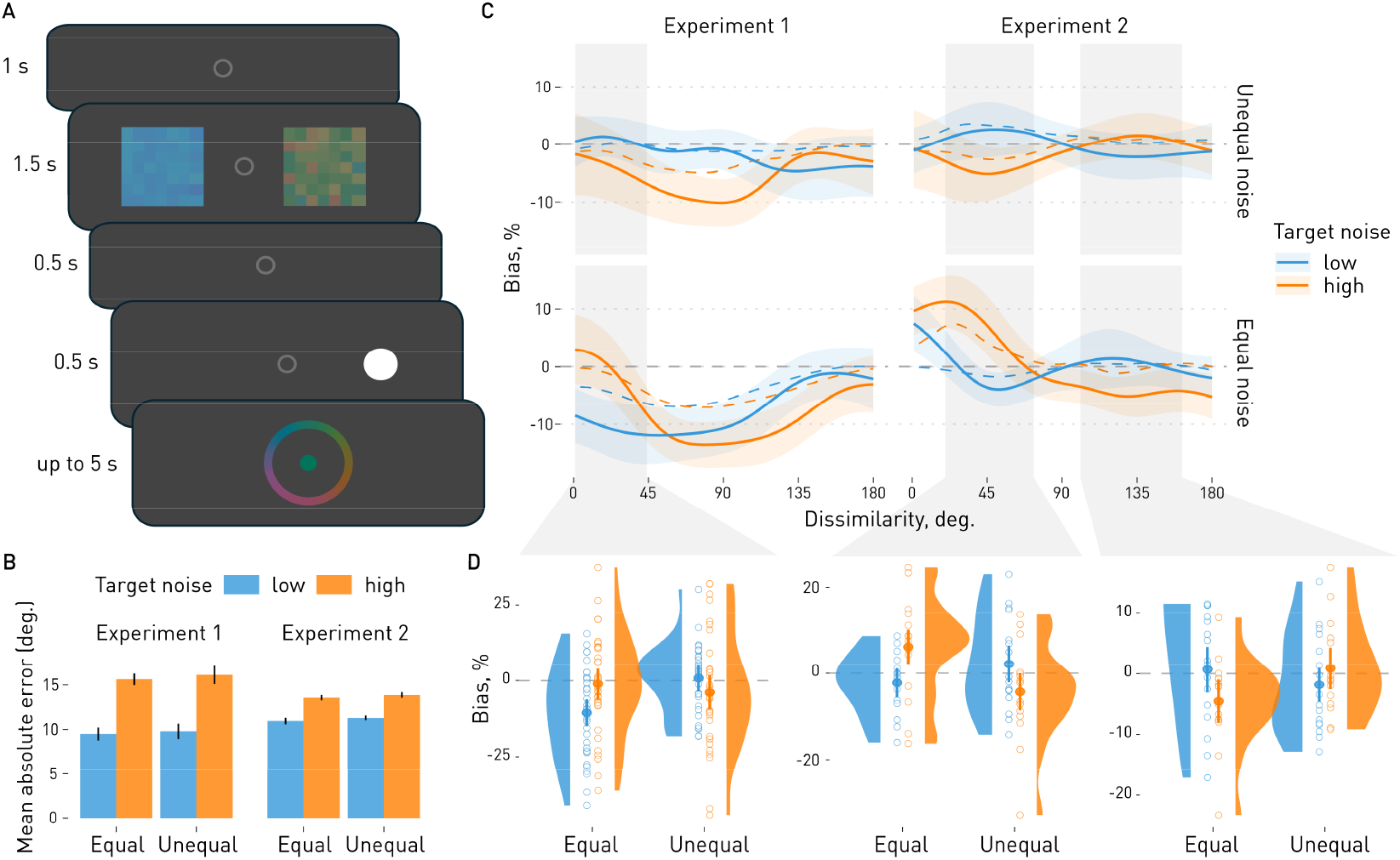
Experimental procedure and results of Experiments 1-2. **A**. All experiments in this paper used the same general procedure. Each trial started with a brief fixation, followed by two mosaic stimuli composed of small squares with varying hues. The hues of each stimulus were drawn from a normal distribution centered on a randomly selected hue. After a short delay, a cue (white circle) indicated which item to report. Participants reported the average hue of the cued item. In Experiments 2–4, cue presentation and report were then repeated for the second item (not shown). **B**. Overall performance in both experiments changed as a function of target noise, whereas non-target noise (and thus the noise equality condition) did not affect error magnitude. Bars show average absolute errors; error bars show 95% confidence intervals. **C-D**. Biases in Experiments 1–2 as a function of target–non-target dissimilarity, target noise level, and the match between target and non-target noise. In the equal noise condition, target and non-target had the same noise level. In the unequal noise condition, non-targets had low noise when targets had high noise, and vice versa. **C**. Continuous bias estimates, with the human data shown as solid lines and model fits as dashed lines; shaded areas show 95% confidence intervals for the mean bias. Gray regions indicate dissimilarity ranges with a significant interaction between target noise and noise equality. **D**. Average biases in the significant clusters in C for each participant (empty circles), their distributions within each condition (slabs), and condition means with 95% confidence intervals (solid circles and bars). Positive bias corresponds to attraction, negative to repulsion.

Observers reported the average color of an item cued with a short delay after the stimuli presentation. The hues of each stimulus were randomly sampled from a wrapped normal distribution, and we manipulated the amount of noise in the stimuli by varying the widths of their distributions in a 2 × 2 design. The reported item (the target) could have low (*SD* = 5°) or high noise (SD = 25° or 45° in Experiment 1, or SD = 25° in Experiment 2; see Methods). The non-target item could have the same amount of noise as the target (‘equal noise’ condition) or a different one (‘unequal noise’ condition, in which case it was the opposite of the target noise, i.e., a high-noise target combined with a low-noise non-target or vice versa). In Experiment 2, following the first color report, observers were additionally cued to report the other item in each trial, in which case the designations of target and non-target were swapped in the data analysis.

#### Overall performance

We first assessed whether the noise manipulation affected response accuracy as a sanity check. Increasing target noise decreased performance in both experiments (Figure 2B; *F*(1, 34) = 91.51, *p* < .001, *η*^2^ _G_= .49 and *F*(1, 15) = 196.23, *p* < .001, *η*^2^ _G_ = .30 for Exp. 1 & 2, respectively). Performance was slightly worse in the unequal noise condition (*F*(1, 34) = 5.24, *p* = .028, *η*^2^ _G_ < .01 and *F*(1, 15) = 6.35, *p* = .024, *η*^2^ _G_ < .01), while the interaction between the factors was not significant (*F*(1, 34) = 0.25, *p* = .617, *η*^2^ _G_ < .01 and *F*(1, 15) = 0.03, *p* = .856, *η*^2^ _G_ < .01). This shows that we were able to manipulate the precision of mnemonic representations as intended, with any effect of noise equality on precision being negligible compared to the manipulation of the target noise.

#### Biases in reproduced colors

We estimated biases based on the dissimilarity between target and non-target colors, target noise level, and noise equality condition. Biases were measured as the difference in probabilities of responding towards versus away from non-targets on a −100% (complete repulsion) to +100% (complete attraction) scale. They were estimated continuously as a function of dissimilarity, with significant effect clusters identified via permutation testing (Figure 2C; see Methods and Figure S2 for details on bias estimation).

In Experiment 1, target noise (from 61° to 118°, *p*_*perm*_ = .013) and noise equality (from 27° to 108°, *p*_*perm*_ = .004) had main effects moderated by a significant interaction (from 1° to 44°, *p*_*perm*_ = .047). In the unequal noise condition, within this dissimilarity range, low-noise targets were numerically more attracted than high-noise targets (*M* = 0.74 [−2.90, 4.38] vs. *M* = −3.93 [−10.39, 2.54], *t*(34) = 1.30, *p* = .202). In contrast, they were repulsed significantly more strongly in the equal noise condition (*M* = −10.58 [−15.54, −5.62] vs. *M* = −1.14 [−6.69, 4.41], *t*(34) = −2.85, *p* = .007; Figure 2C-D).

In Experiment 2, we found a main effect of noise equality (from 1° to 45°, *p*_*perm*_ = .037), but crucially also two clusters with significant interaction between target noise and noise equality (from 20° to 73°, *p*_*perm*_ = .020 and from 101° to 162°, *p*_*perm*_ = .010). In the first cluster with intermediate color dissimilarity, low noise targets were attracted more than the high noise ones under unequal noise (*M* = 2.12 [−3.70, 7.95] vs. *M* = −4.33 [−10.36, 1.69], *t*(15) = 1.97, *p* = .067) but repulsed more when noise levels were equal (*M* = −2.26 [−5.94, 1.42] vs. *M* = 6.09 [−0.03, 12.21], *t*(15) = −3.00, *p* = .009). In the second cluster at larger dissimilarity, the effect was in the opposite direction with numerically less attraction for low noise targets under unequal noise (*M* = −1.85 [−6.18, 2.48] vs. *M* = 0.87 [−3.09, 4.83], *t*(15) = −1.68, *p* = .114), but more attraction in the equal noise condition (*M* = 0.68 [−4.28, 5.64] vs. *M* = −4.54 [−8.36, −0.71], *t*(15) = 2.33, *p* = .034). We also repeated this analysis including the response order (first vs. second response). While the second response showed stronger attraction, the interaction between the noise equality and target noise was visible for both responses and no three-way interaction was observed (Figure S3).

#### Can the model fit the data?

We then tested whether the Demixing Model could reproduce the nonlinear patterns observed in the empirical data (Figure 2C). The model predictions were simulated for a dense parameter grid (noise and similarity in the color dimension; discriminability in the identifying dimension; see Methods for details) and interpolated with a 4-layer feedforward neural network to enable fast estimation of model likelihoods. The fit between the model predictions and each participant’s data was then optimized using a loss function combining the correlation and the squared distance between the predicted and empirical bias curves. The noise parameters for the color dimensions were set free for each of the four conditions (target noise × noise equality), while the noise on the identifying dimension was assumed to be fixed for each participant. The model achieved a strong correlation with the empirical bias data (*r*_pearson_: *M* = 0.91, *SD* = 0.08, range = [0.59, 0.99] across conditions and observers). Thus, despite having only two free noise parameters in each condition, the model could approximate the non-linear patterns of human biases.

### Experiment 3

Experiments 1 and 2 supported the predictions of the Demixing Model, demonstrating an interaction between the target item noise level and the noise equality condition. As hypothesized, low-noise targets were attracted more than high-noise targets in the unequal noise condition, but repelled more in the equal noise condition. In Experiment 3, we aimed to replicate the main findings of Experiments 1 and 2. Instead of varying dissimilarity across the whole 180-degree range, we focused on three specific points corresponding to the clusters where the interaction between target noise and noise equality was observed in these experiments. The dissimilarity between target and non-target was set to 20°, 45°, or 135°, with ±3° variation.

#### Overall performance

In Experiment 3, we again observed a significant effect of target noise on recall accuracy (*F*(1, 22) = 127.53, *p* < .001, *η*^2^ _G_ = .09) but no effect of noise equality (*F*(1, 22) = 2.69, *p* = .115, *η*^2^ _G_ < .01) or their interaction (*F*(1, 22) < 0.01, *p* = .952, *η*^2^ _G_ < .01). The performance was better for low-noise (M = 12.21 [10.85, 13.96]°) than for high-noise targets (M = 14.79 [13.30, 16.51]°).

#### Biases in reproduced colors

The biases were affected by an interaction between target noise level and noise equality condition (*F*(1, 22) = 29.49, *p* < .001, *η*^2^_G_ = .03), further qualified by a three-way interaction with similarity bin (*F*(2, 42) = 3.86, *p* = .031, *η*^2^_G_ < .01; Figure 3). The target noise and noise equality interaction was significant at 20° (*F*(1, 22) = 19.52, *p* < .001, *η*^2^_G_ = .06) and 45° (*F*(1, 22) = 9.69, *p* = .005, *η*^2^_G_ = .03) dissimilarity, but not at 135° (*F*(1, 22) = 1.29, *p* = .269, *η*^2^_G_ < .01). The additional analyses with the response order factor included showed the same pattern of results. In addition, we found a stronger attraction in the second response but no interaction with other factors (Figure S3).

**Figure 3.**
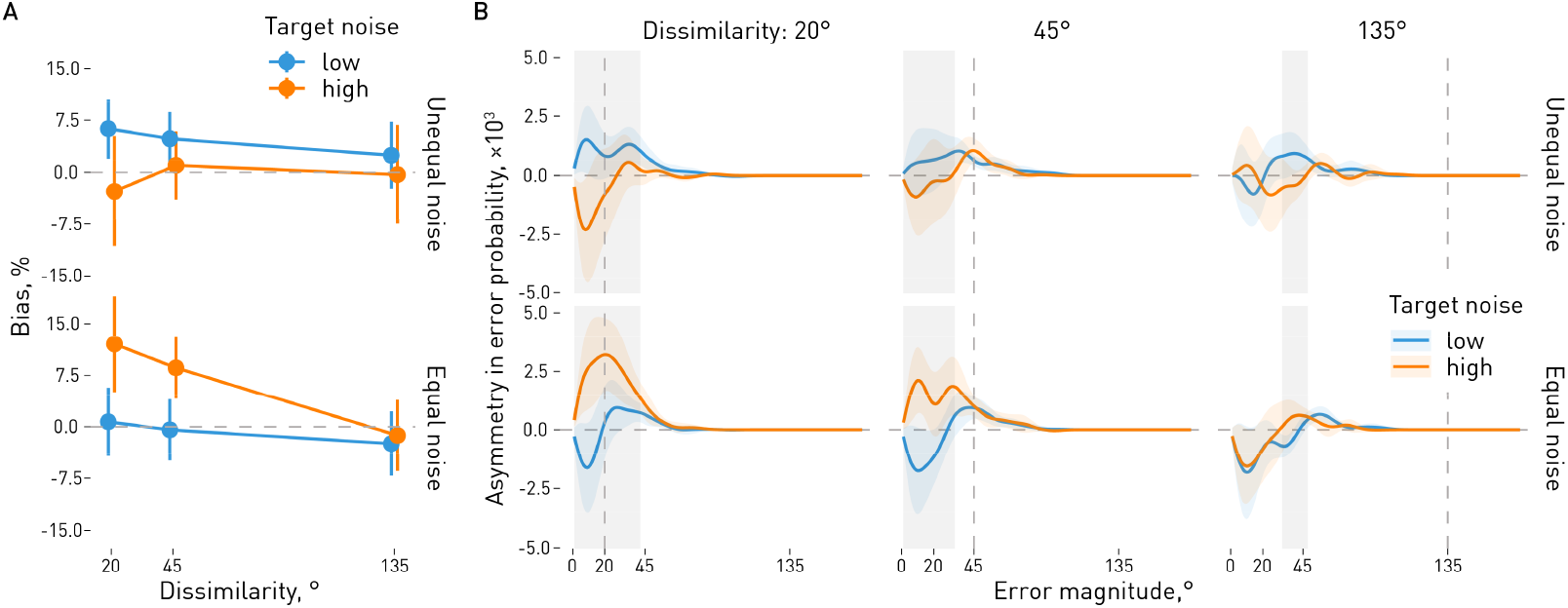
Biases in Experiment 3 as a function of dissimilarity and the noise levels. **A**. Average biases. In the equal noise condition, target and distractor had the same noise level. In the unequal noise condition, distractors had low noise when targets had high noise, and vice versa. Replicating previous results, in the unequal noise condition low-noise targets were more strongly attracted than high-noise targets, whereas in the equal noise condition the opposite pattern was observed. **B**. Error probabilities as a function of error magnitude. To test whether attractive biases were driven by swap errors, we computed an asymmetry index (difference in probability of attractive minus repulsive errors, y-axis) for each error magnitude (x-axis). That is, for each possible error magnitude (the absolute value of response deviation from the stimulus), we measured how much more likely responses were to fall toward the non-target than away from it. The interaction between target noise and noise equality is already present for relatively small errors, which is inconsistent with swap errors being the main driving factor. Positive bias and asymmetry index correspond to attraction, negative to repulsion. Lines show mean asymmetry with shaded areas indicating 95% confidence intervals; gray regions indicate dissimilarity ranges with a significant interaction between target noise and noise equality.

In the 20° bin, low-noise targets were more attracted towards non-targets under unequal noise (*M* = 6.25 [0.95, 11.55] vs. *M* = −2.73 [−13.67, 8.21], *t*(22) = 2.21, *p* = .038). In contrast, high-noise targets were attracted more under equal noise (*M* = 0.71 [−7.59, 9.00] vs. *M* = 12.01 [2.80, 21.22], *t*(22) = −3.74, *p* = .001). The same reversal occurred in the 45° bin, where low-noise targets were numerically more attracted under unequal noise (*M* = 4.82 [−1.34, 10.97] vs. *M* = 1.00 [−7.87, 9.86], *t*(22) = 1.21, *p* = .238), but high-noise targets under equal noise (*M* = −0.46 [−7.86, 6.94] vs. *M* = 8.58 [0.96, 16.20], *t*(22) = −3.13, *p* = .005).

Are attractive biases observed for low-noise targets in the unequal noise and for high-noise targets in the equal noise condition explained by swap errors? To answer this question, we computed the error asymmetry as a function of the error magnitude. The asymmetry was computed as a difference in probability of errors towards the non-target item (attractive errors) and away from it (repulsive errors). If swap errors were driving the observed effects, we would expect that the asymmetry in error probabilities would be affected most for errors clustered around the non-target color. Instead, we found that the crucial interaction between the target noise and noise equality was present already for relatively small errors, unlikely to be caused by swaps with non-targets. In the 20° bin, the interaction effect was significant already from 1° to 42°, *p*_*perm*_ < .001; in the 45° bin, it was significant for errors from 1° to 33°, *p*_*perm*_ < .001. Interestingly, it was even observed in the 135° bin for errors between 32° and 48°, *p*_*perm*_ = .032, even though this effect was invisible in the aggregate data analysis above. In sum, the asymmetry in error probabilities is present already for relatively small errors, unlikely to be caused by swaps with non-targets (see also Figure S4 for the analysis of small errors only).

### Experiment 4

Experiment 3 replicated the findings of Experiments 1 and 2, clearly showing attractive biases for low-noise targets in the unequal noise condition and repulsive biases for high-noise targets in the equal noise condition. Together with the previous experiments, the results support the Demixing Model predictions. However, in the unequal noise condition of these experiments, a change in the relative level of target noise—whether it is higher or lower than the non-target noise—also resulted in a change in the absolute level of target noise. That is, when the target became noisier than the non-target, this was achieved by adjusting the target noise level simultaneously with the change in the non-target noise level.

In Experiment 4, we tested whether changes in relative noise levels affect biases when the absolute target noise level is held constant, as predicted in Figure 1D. The task and general procedure were identical to Experiment 3, except that the target item always had the same noise level (*SD* = 20°). To manipulate relative noise, we varied only the non-target noise level, making the target noisier than the non-target (non-target *SD* = 5°), equally noisy (non-target *SD* = 20°), or less noisy (non-target *SD* = 35°). In line with Experiment 3, the participants gave responses to both items on each trial; however, in the analyses of biases we focus mostly on the target with the constant noise level in line with the predictions of the model.

#### Overall performance

The relative level of target noise did not affect the recall accuracy (measured as the mean absolute error) when the absolute target noise level was kept constant (*F*(2, 77) = 0.49, *p* = .609, *η*^2^_G_ < .01). In contrast, when recalling the item with varying noise levels, the effect of noise level was significant as expected (*F*(2, 64) = 58.66, *p* < .001, *η*^2^_G_ = .13), with larger errors for noisier targets.

#### Biases in reproduced colors

We found a significant effect of the relative target noise level on the bias magnitude and direction (*F*(2, 67) = 7.51, *p* = .002, *η*^2^_G_ = .03), further qualified by an interaction with the similarity bin (*F*(4, 139) = 13.04, *p* < .001, *η*^2^_G_ = .06). Further analyses showed that the effect of relative target noise was significant in the 20° similarity bin (*F*(2, 76) = 21.81, *p* < .001, *η*^2^_G_ = .15) and in the 45° similarity bin (*F*(2, 68) = 9.54, *p* < .001, *η*^2^_G_ = .06), but not in the 135° similarity bin (*F*(2, 65) = 1.95, *p* = .157, *η*^2^_G_ = .02).

As Figure 4A shows, in the 20° bin, the repulsion was stronger for targets with higher noise level (*M* = −15.93 [−21.84, −10.01]) compared to the same (*M* = 1.75 [−2.31, 5.81]) or lower (*M* = 2.12 [−3.06, 7.30]) noise. In the 45° bin, in contrast, the targets with lower noise were attracted more strongly (*M* = 6.10 [1.57, 10.63]) than the targets with the same noise (*M* = −2.81 [−5.61, −0.00]) or higher noise (*M* = −4.17 [−8.72, 0.37]). And as indicated by the ANOVA results described above, in the 135° bin, there was no significant difference between the relative noise levels (*M* = −1.17 [−4.60, 2.26] for same, *M* = 2.86 [−2.82, 8.54] for higher, and *M* = −3.68 [−10.59, 3.23] for lower noise).

**Figure 4.**
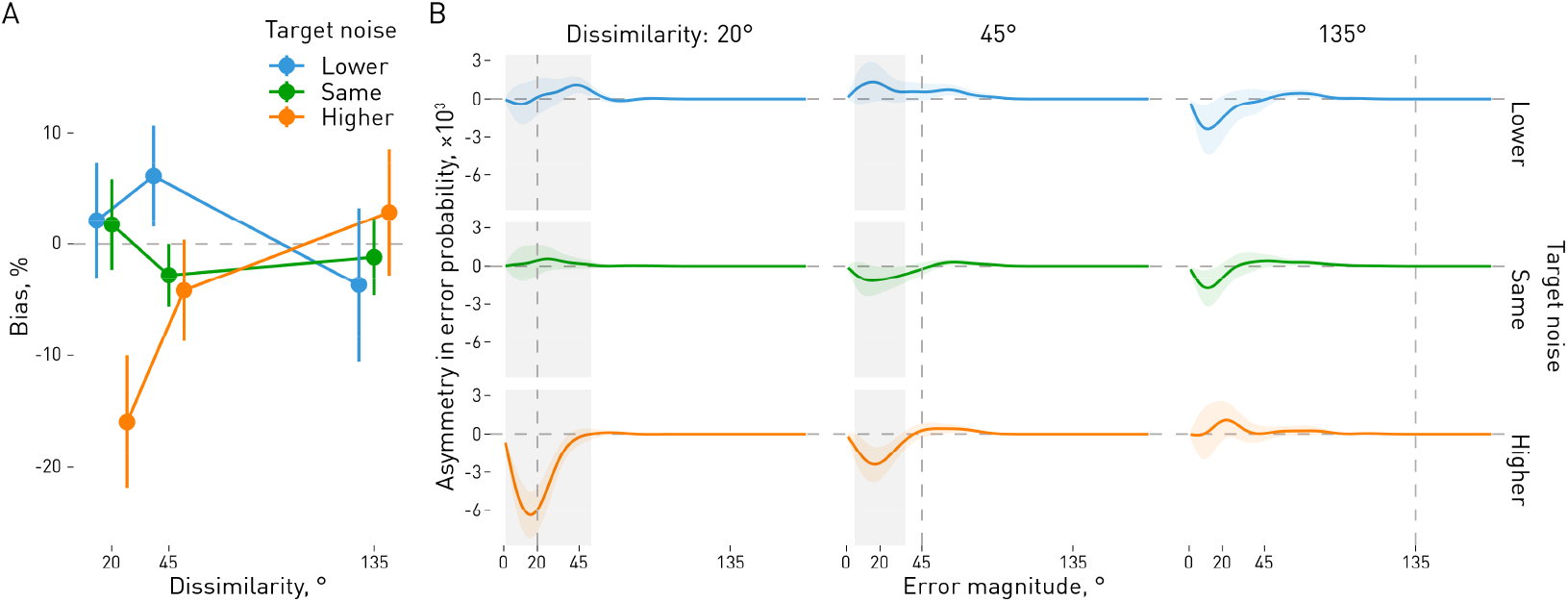
Biases in Experiment 4 as a function of dissimilarity and the relative noise levels. **A**. Average biases. Target noise level was kept constant while non-target noise varied, so the target could have higher, equal, or lower noise than the non-target. In line with the model predictions, targets were repulsed more when they had higher noise than the non-target and attracted more when they had lower noise. Points show mean biases with 95% confidence intervals (bars). **B**. As in Experiment 3, we estimated an asymmetry index in error probability (y-axis) for errors of different magnitudes (x-axis). That is, for each possible error magnitude (the absolute deviation of the response from the target), we calculated the difference in the probability of errors toward the non-target versus away from it. In the 45° bin, which showed the largest attractive bias in the lower-noise condition, the effect of target noise is already present for relatively small errors, inconsistent with swap errors being the main driving factor of the differences between conditions. Positive bias and asymmetry index correspond to attraction, negative to repulsion. Lines show mean error asymmetry with shaded areas indicating 95% confidence intervals; gray regions indicate clusters with a significant effect of target noise.

We then analyzed the asymmetry in error probabilities as a function of error magnitude separately for each similarity bin, following the same procedure as in Experiment 3. The relative target noise effect was significant already for relatively small errors in the 20° bin (from 1° to 52°, *p*_*perm*_ < .001), and crucially, in the 45° bin (from 5° to 35°, *p*_*perm*_ = .011) that showed the significant attractive bias in the average bias analyses in the lower noise condition. In the 135° bin, no significant clusters were observed even though the results look qualitatively similar to Experiment 3. This suggests that while swap errors might play a role in biases, the attraction observed for the lower noise condition stems from relatively small errors, reflecting a drift in representations.

## Discussion

Why are working memory reports biased when multiple items are held in memory? The Demixing Model^22^ offers a principled answer: when signals from multiple items overlap, disentangling them inevitably produces biased estimates. Crucially, the model predicts that these biases depend not only on the overall level of noise, but also on how noise is distributed between the memorized items. The less noisy item should show more attraction (less repulsion), whereas the noisier one should show more repulsion (less attraction), compared to when both items have the same noise level (i.e., both are low-noise or both are high-noise).

Our experiments tested these predictions using a novel noise manipulation that varied the color variability of memorized items. In Experiments 1 and 2, the bias direction depended on the relative noise of target and non-target items: with equal noise, low-noise targets were repelled more (attracted less) than high-noise targets, whereas with unequal noise they were attracted more (repelled less) than high-noise ones. Experiment 3 replicated this asymmetry while focusing on the similarity levels (20°, 45°, and 135°) with the strongest effects in previous experiments. The same reversal between equal- and unequal-noise conditions remained, even when we restricted analyses to small errors to rule out swap explanations. Finally, Experiment 4 held target noise constant while varying non-target noise, still finding that relative noise dictated whether targets were more attracted or repelled more. Across all four experiments, results consistently aligned with the counterintuitive predictions of the Demixing Model.

Our results build on previous studies showing that memory noise is important in determining the direction of biases ^5,9^. However, we show that it matters not just how much noise there is in total, but also how it is distributed between the target and non-target items. Increasing the amount of noise while keeping it equally distributed between the stimuli leads to effects that are qualitatively different from those observed when only target noise or only non-target noise is increased. The *relative* noise level is as important as the *absolute* one.

Attractive biases in memory are often attributed to confusion or integration between items ^5,25,26^. In our equal-noise condition, noisier targets were indeed attracted more toward non-targets than less noisy targets, consistent with this account. However, if this simple explanation were correct, we would also see the same pattern in the unequal noise condition in Experiments 1-3 (or no effect, as the overall noise amount was the same regardless of the target noise). The finding of more attraction for low-noise targets under unequal noise is inconsistent with a simple confusion account and instead points to relative noise as the critical determinant of bias direction.

Repulsive biases have been attributed to the need to reduce interference between items in memory by pushing them further apart ^5,21^. Chunharas and colleagues^5^ further suggested that “repulsion is expected whenever items are similar enough, and uncertainty high enough” (p. 15). Our data support the first but not the second part of this prediction. The biases (all biases, not just repulsion) indeed decrease if the dissimilarity is very high, although the overall pattern is non-linear. However, higher uncertainty does not necessarily lead to stronger repulsion. When the noise levels between the target and non-target are kept the same, increasing noise decreases repulsion (increased attraction) for similar targets. Furthermore, in the unequal noise condition, the overall noise levels stay the same whether the target is assigned a low or a high noise level, yet we still see the change in the bias strength with more repulsion (less attraction) for high noise targets. These changes are incompatible with the idea of “interference reduction” as the computational goal behind the biases, a conclusion resonating with the recent work showing that serial biases do not reduce response variability^27^.

Repulsive biases are also often explained with efficient coding in the case of sequentially shown stimuli^28–31^ or biases relative to references, such as cardinal axes in orientation perception^32–35^. Efficient-coding models posit that the brain allocates neural resources to reflect environmental statistics, predicting repulsion from the most likely stimulus values, whether from prior trials or those prevalent in nature. However, this framework struggles to account for biases among simultaneously presented items, as the brain must first identify which stimuli to prioritize or suppress, a challenge when stimuli are shown together. While efficient coding may explain other kinds of biases, it is difficult to see how it can account for our findings.

We based our study on the predictions of the Demixing Model, and supporting these predictions lends support to the model. Furthermore, we demonstrated that the Demixing Model fits data well, aligning with patterns observed in human observers, despite being a normative model with only a few free parameters for stimulus noise. However, as Figure 2C illustrates, the model struggled to capture certain data patterns (e.g., an attractive bias followed by a repulsive one as dissimilarity increases). Some discrepancies can be attributed to factors not considered in our current analysis, such as the influence of a trial’s first response on subsequent responses. While we focused on testing specific model predictions rather than providing post hoc explanations, offering a more comprehensive account is essential for advancing computational and theoretical development.

Our results also reveal that memory biases follow strikingly non-linear dynamics as a function of item similarity. Across Experiments 1 and 2, biases did not simply become more attractive or more repulsive with increasing dissimilarity but changed in non-monotonic ways. Interestingly, previous studies showed both attraction followed by repulsion ^5^ and repulsion followed by attraction ^2,4,36^ but our results show that both are possible for simultaneously shown stimuli differing only in noisiness. This means that focusing on a single similarity level could easily lead to misleading conclusions or even mask the influence of noise altogether. More generally, while restricting the number of similarity levels can increase statistical power, our data show that it is crucial to first map out the full similarity space before targeting specific points within it.

While our study focuses on visual working memory, similar explanations based on disentanglement of noisy signals can be applied to other contextual effects, from perceptual, such as the tilt illusion ^37,38^ or serial dependence ^39–41^, to high-level ones, such as the decoy effects ^42–44^. However, potential changes in representation format^45,46^ and other temporal dynamics^2,36^ require extra caution when interpreting the effects of noise in sequential effects, such as serial dependence. Similarly, the effects of noise in high-level decisions constitute a very promising avenue of study^44,47^, but manipulating noise in subjective values requires extra caution, as it is not directly controllable by the experimenter. From this perspective, simple visual stimuli, like the color patterns here, might be an ideal model system for understanding basic mechanisms of contextual biases, laying the groundwork for further research in other domains.

In sum, our results highlight the central role of noise in determining the direction and magnitude of biases in visual working memory. The asymmetry in biases for low- and high-noise items, as well as the changes in biases due to non-target noise, challenge dominant accounts that attribute attraction merely to confusion or repulsion to interference reduction. Instead, the findings support the view that biases arise from the need to disentangle overlapping signals from their corresponding sources, as formalized by the Demixing Model. From this perspective, memory reports are not just corrupted traces of isolated items, nor merely interactions between them, but reconstructions that are inevitably affected by inter-item structure. Visual working memory thus fits within the broader framework of reconstructive memory: biases are not anomalies but signatures of the processes that make memory possible.

## Methods

### Experiments 1 & 2

#### Participants

Experiment 1 consisted of two samples (Exp. 1A/1B, see details below). 18 participants took part in Exp. 1A, recruited via Prolific based on their performance in a short 5-minute version of the task from a larger sample of 43 participants. The top eighteen performers were invited to the main study. Participants were paid £1.20 for the pre-selection phase and £6 for the main study. This and the later studies were approved by the Board for Research and Research Training (FFU) at the Faculty of Psychology, University of Bergen. Participants provided electronic informed consent before the study began. Age and gender data were not recorded, as we did not aim to analyze these variables and wished to avoid collecting personal data in compliance with local regulations. Seventeen participants took part in Exp. 1B, recruited at the University of Iceland. 16 participants were recruited via Prolific for Experiment 2 with the same compensation. The minimal sample size was determined based on the simulations that indicated that nine participants should be enough to detect a 5% difference in biases with the design we used. However, we based our sample sizes on a more conservative estimate of at least 16 people.

#### Materials and procedure

Participants performed a color visual working memory task with stimuli varying in hue with different amounts of noise added. In Experiment 1, they saw a post-cue telling which item to report, while in Experiment 2 they were also required to report the second item.

The experiments were conducted online using jsPsych (v.7.3.1, de Leeuw, 2015) with the psychophysics plugin (v.3.5.0, Kuroki, 2021) and administered through JATOS (Lange et al., 2015). They required a minimum resolution of 1000 × 600 pixels, and participants were instructed to switch to a full-screen mode.

Trials began with a fixation circle (25 px radius; 1000 ms), followed by two colored patches (1500 ms) on the left and right (±300 px) against a neutral background (25% luminance). Each patch consisted of an 8 × 8 grid of 32 × 32 px colored squares varying in hue (OKLCH color space: luminance 50%, chroma 0.1). Participants were asked to remember them and later report the average color of one (Exp. 1) or both (Exp. 2) patches. The hues were normally distributed with random means for each patch and the SD of 5° for low-noise and 20° (Exp. 1A, Exp. 2) or 45° (Exp. 1B) for the high-noise items. Since the results from Exp. 1A and 1B were similar, they are presented together.

After a 500 ms delay, a white cue appeared at either the left or right position for 500 ms, indicating which patch to report. Participants then selected the color that best matched the cued patch by clicking on a color wheel (150 px radius, 30 px thickness, random hue offset) within 5000 ms. In Experiment 2, participants were then presented with a second cue at the opposite location and reported the second patch color using the same procedure. Finally, they got feedback in the form of a numerical score based on their accuracy.

The experiment began with a training phase of 10 trials during which presentation times were initially longer (4000 ms for patches; 2000 ms for cues) and gradually decreased to match the timing in the main part. Participants received feedback with their response and the correct color during training.

Participants did two (Experiment 1) or four (Experiment 2) sessions with 304 trials split across two blocks within each session with target position (left, right), target standard deviation (5°, 45°), and noise equality (equal noise for two items vs. unequal noise) counterbalanced within a block. Since Experiment 2 required two reports, in the following we refer to the item cued before each response as the target.

#### Data preprocessing

We used the *circhelp* package in *R* (Chetverikov, 2023) to remove anisotropy in color reports and exclude outliers. In brief, for each participant, we estimated the mean and variability of errors as a function of color, while allowing for discontinuity around one or more colors to account for potential repulsive biases from category boundaries. Responses that deviated by more than three standard deviations from the mean error at a given color were considered outliers and excluded from further analyses. The anisotropies were then removed by subtracting the mean color-dependent error from each response. A more detailed description of this procedure can be found in the package documentation.

#### Data analysis

If observers are biased, they would more often make errors in the direction of a non-target item (attractive bias) or away from it (repulsive bias). More technically, biased responses are characterized by the asymmetry of the probability distribution of responses around the true stimulus feature value. To quantify this asymmetry, we first recoded response errors in such a way that the positive sign corresponds to errors in the direction of non-targets, while a negative error corresponds to errors away from them (Figure S2A). We then computed the probability distribution density for errors as a function of similarity between targets and non-targets across all trials. In Exps. 1-2, where the similarity was set randomly on each trial from the full range of available values (180 degrees for color), we used a weighted kernel density estimation approach. For each possible similarity value *i* across the feature space discretized in 1-degree steps, we determined the weight for a given trial using the Gaussian kernel as *w*_*i*_ =*N*(*i*, |*s*_*t*_ −*s*_*d*_|,*σ*_*f*_), where *s*_*t*_ is the target stimulus, *s*_*nt*_ is the non-target, | *s*_*t*_ - *s*_*nt*_ | denotes the absolute circular difference (dissimilarity) in their values, *σ*_*f*_ is a feature-dependent kernel width (20 degrees; other widths provide similar results). In Exps. 3 and 4, where only some bins of the full similarity space were used, a simple probability density was computed for each similarity bin.

We then computed the bias as the asymmetry of the error distribution: *B* = ∑*P*_+_ - ∑*P*_−_, that is, the sum of probability density for positive errors minus the sum of probability density for negative errors, and converted the result to a scale from −100 to 100 percent. In other words, our bias measure reflects how much more likely it is for observers to err towards than away from them. A value of zero indicates no bias, a value of 100% indicates that observers always err towards the non-target (attractive bias), and a value of −100% indicates that they always err away from the non-target (repulsive bias).

The significance of conditions’ effects on bias magnitude and direction, given biases that vary with dissimilarity, was assessed using a permutation-based procedure. First, a within-subject ANOVA was conducted for each 1-degree dissimilarity step. Next, clusters of significant effects were identified based on the *F* value corresponding to a conventional *p* < .05 criterion, which is solely for cluster identification and does not imply sharp boundaries. We then calculated the likelihood of obtaining a cluster of this or larger size by chance. In 10000 replications, we shuffled the condition labels and repeated the procedure, measuring the cluster sizes for each effect of interest (e.g., target noise, noise equality condition, or their interaction). Finally, *p*-values were computed as the probability of observing a cluster of a given size across permutations.

### Experiment 3

#### Participants

Twenty-four participants took part in Experiment 3. They were either the students (*N* = 10), who took part in exchange for the course credits, or were recruited via Prolific (*N* = 14), in which case they received monetary compensation with the same per-hour rate as in other studies. One participant was excluded because their response variability was very high (*SD* = 84°).

#### Procedure

Experiment 3 followed the design of Experiment 2 with the exception that the similarity of the two stimuli in each trial was picked randomly from one of three bins with a ±3° range centered at 20°, 45°, and 135°. Each participant took part in 2 sessions with 596 trials each (two participants completed one session only due to technical issues).

#### Data analysis

In Experiments 3 and 4 we additionally analyzed the asymmetry in error probabilities for errors of different magnitude, computed as the difference in probability of errors towards the non-target item (attractive errors) and away from it (repulsive errors) as a function of the error magnitude. For each participant in each combination of conditions, we estimated the error probability density (indicating how likely the errors of different magnitudes are) on a −180 to 180° grid with a 1° step and subtracted the negative side corresponding to repulsive errors from the positive side corresponding to attractive errors. We then used repeated measures ANOVA to test the effects of target noise and noise equality on the asymmetry in error probabilities for each 1° step of error magnitude. Then, we extracted clusters of significant interaction effects. Finally, we used permutation testing identical to the ones used in other cluster-based analyses in Exp. 1 and 2 to assess the probability of observing clusters with the same or larger size due to chance.

### Experiment 4

#### Participants

Forty-two participants took part in the study, recruited via Prolific (*N* = 22) or from a student population as part of the course projects (*N* = 20). They were compensated at the same per-hour rate as in previous studies. Two were excluded due to high response variability (*SD* = 62° and 71°).

#### Procedure

Experiment 4 followed the design of Experiment 3, with the exception that in each trial, the noise level of one item (the ‘target’) was kept constant at *SD* = 20°, while the noise level of the other item was manipulated to be either lower (*SD* = 5°), the same (*SD* = 20°), or higher (*SD* = 35°) than the first item’s noise. The observers again reported both items following the same procedure as in previous experiments. Each participant took part in a single session consisting of 576 trials in the main part and 10 training trials.

### Simulations and model fits

We first simulated the probabilities of biases of different magnitudes (their likelihood surfaces) as a function of similarity between the target and non-target items on a 20 × 20 × 20 grid of the model parameters (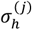and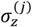) in a 10–200 range (in ° for hue or unitless for identifiability). The noise levels of the target and non-target items 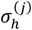 varied independently while the noise level in the identifiability dimension 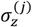 was assumed to be the same for both items (so 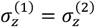). The mean values of the stimuli in the hue domain 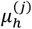 were set around the 0° point with dissimilarity between them set on a grid from 4° to 180° with 4° steps (e.g., 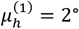 and 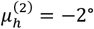 for a dissimilarity of 4°; since hue is circular, the choice of the starting point does not matter). The means in the identifiability dimension 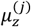 were fixed at ±24° since 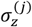 and 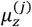 variation could be used interchangeably to achieve the same effects in terms of how well the items can be discriminated (that is, discriminability is a ratio of the difference between the means and their standard deviation).

For each of the parameter combinations, we sampled 10000 data samples from a bivariate wrapped normal – normal distribution (Eq. 1). Each sample had *N* = 20 observations (*N* = 100 yielded qualitatively similar results). We then used the standard expectation maximization (EM) algorithm with a modified Gaussian mixture model that accounted for circularity in one of the dimensions to obtain the most likely solution for each sample. In brief, EM iteratively searches for the most likely solution by estimating the probability that each observation comes from each stimulus under current parameter values, and then updating those parameters based on the estimates. To ensure the EM convergence, we used 50 random initial starting points for each sample and strict convergence criteria (a relative change of 0.1% or less for the likelihood of the data under the parameters as estimated via the variational lower bound, and the requirement of all 50 parallel estimations to reach both the convergence criteria and the global likelihood maximum, with a limit of 2000 iterations). The minimum standard deviation was set to 5 (in ° for hue or unitless for identifiability) and the weights *σ*_*j*_ were limited to the [0.1, 0.9] range.

We then computed the biases probability for each parameter combination using circular kernel density estimation on a grid of −180 to 180° with 2° steps (181 steps in total). These probability distributions were then combined into log-likelihood surfaces showing the probabilities of biases as a function of bias magnitude and direction and the dissimilarity between stimuli (4° to 180° with 4° steps, 89 steps in total). These log-likelihood surfaces were used as a basis for model predictions in Figure 1, with asymmetry in probability density computed the same way as for the real data as a measure of bias for the ease of comparison with the data.

A neural network was then trained to predict the likelihood surface from the model parameters for fitting purposes, allowing us to interpolate between the points of the parameter grid without simulations. The network had a 4-layer feedforward architecture (256, 512, 1024, and 181×89 = 16109 units, respectively) with 3 parameters as inputs 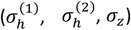 and 181×89 log-likelihood surfaces as outputs. The network was trained for 2,500 epochs using AdamW optimizer with warmup-cosine scheduling (peak learning rate 2×10^-3^, weight decay 1×10^-4^), batch sizes 32-64, and a composite loss function combining MSE, KL divergence, expectation loss, and smoothness regularization (with weights 0.3, 0.3, 0.2, and 0.2, respectively).

To fit the model to the data, we optimized the model parameters for each subject using the density asymmetry curve in each condition as a target. The hue noise parameters 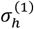 and 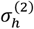 were optimized for each condition separately, while the identifiability noise *σ*_*z*_ was shared across conditions for a given subject. We used a hierarchical grid search over the shared spatial parameter using 20-point initial grids that refine with 0.5 zoom factors until 1° minimum step size. For each shared parameter value, hue noise parameters were optimized using a hierarchical grid search with grid steps decreasing in sequence (10°, 6°, 4°, 2°, 1°). The loss function combines MSE, normalized by dividing by the empirical bias curve range, and correlation loss (1 – Pearson *r*) between predicted and empirical asymmetry curves with weights of 0.75 and 0.25, respectively.

## Data and code availability

Data and code for behavioral experiments and data analyses are available at OSF: https://osf.io/kqb8t/ Model simulations and fitting code is available on Github: https://github.com/achetverikov/demixing_model

## Supplementary Figures

**Figure S1.**
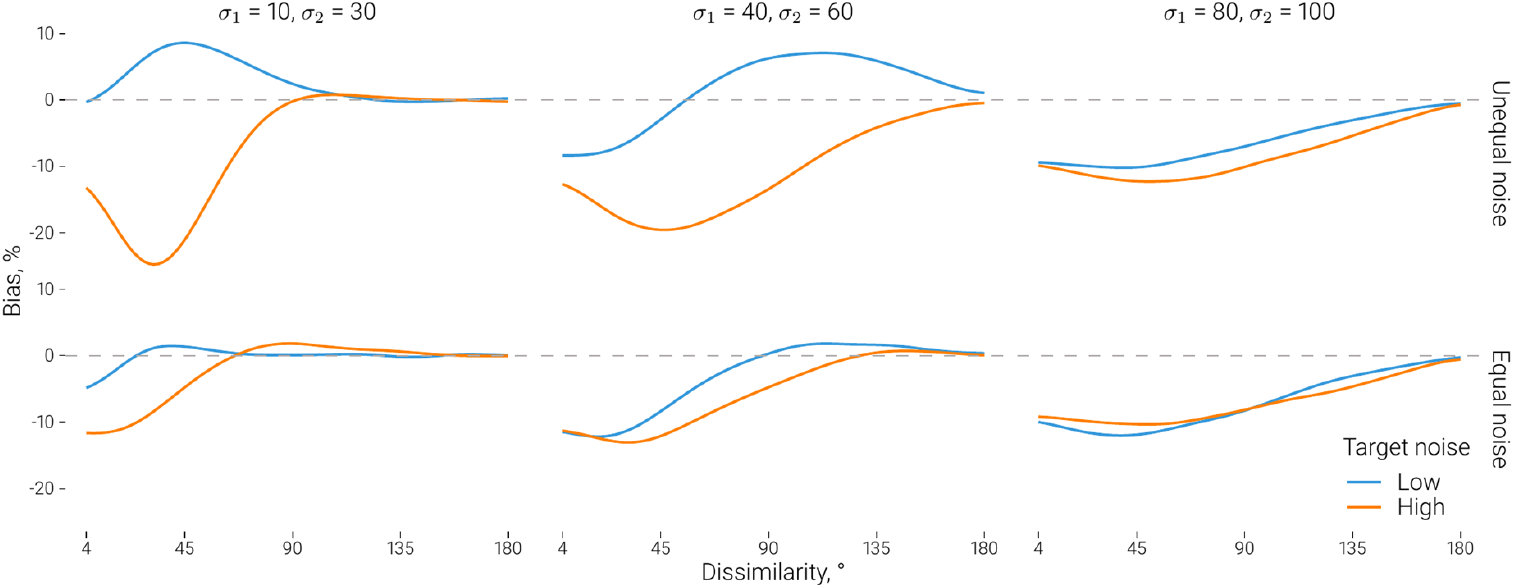
Predictions of the Demixing Model for varying levels of noise for the mixture components. Each column shows the average bias for the case when stimuli noise in the unequal noise case corresponds to the values shown at the top. In the equal noise case, the two components have equal noise levels corresponding to one of these two values. Color corresponds to the target noise level, which is either the same (‘equal noise’) or different (‘unequal noise’) from the non-target.

**Figure S2.**
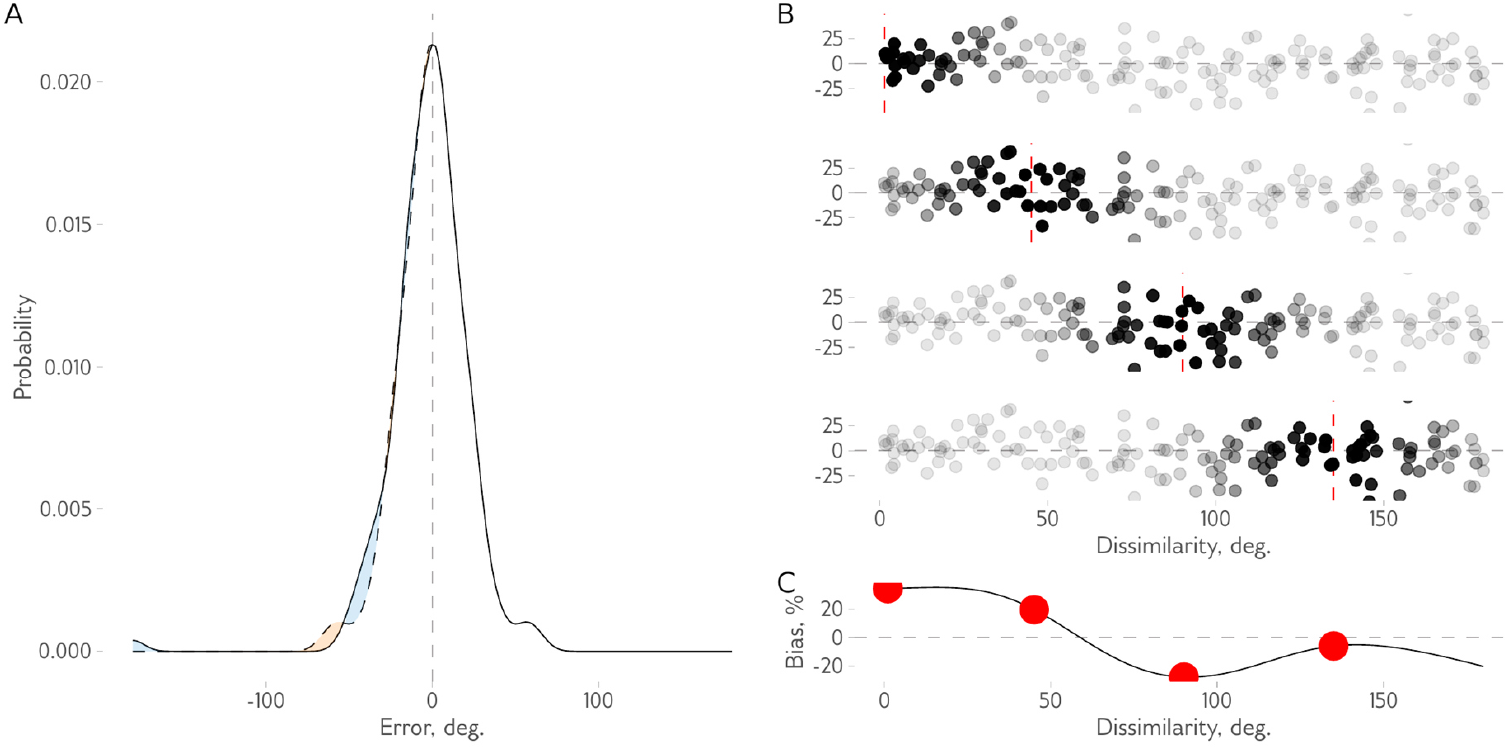
Example bias estimating procedure for a single participant in a single condition. **A** The response probability density (solid line) in the case of biased responses is often asymmetric. This asymmetry is clearly visible when comparing a symmetric version of the same distribution computed by mirroring the positive side of the distribution to the negative values (dashed line). The difference between the two distributions is shown with a shaded area. **B-C** When dissimilarity between target and non-target varies, it might affect the strength and direction of biases. To obtain a continuous estimate of bias as a function of dissimilarity, we computed asymmetry in response probabilities for each 1° step of dissimilarity using weighted probability density estimates, highlighted here for four arbitrary points (1, 25, 50, and 75°). The weights were computed as a Gaussian distribution with the mean at the dissimilarity point and a standard deviation of 20°. **B**. The weights of individual response errors (points), with more transparent points indicating lower weights. **C**. The resulting bias curve as a function of dissimilarity, with 4 red points corresponding to four panels in B.

**Figure S3.**
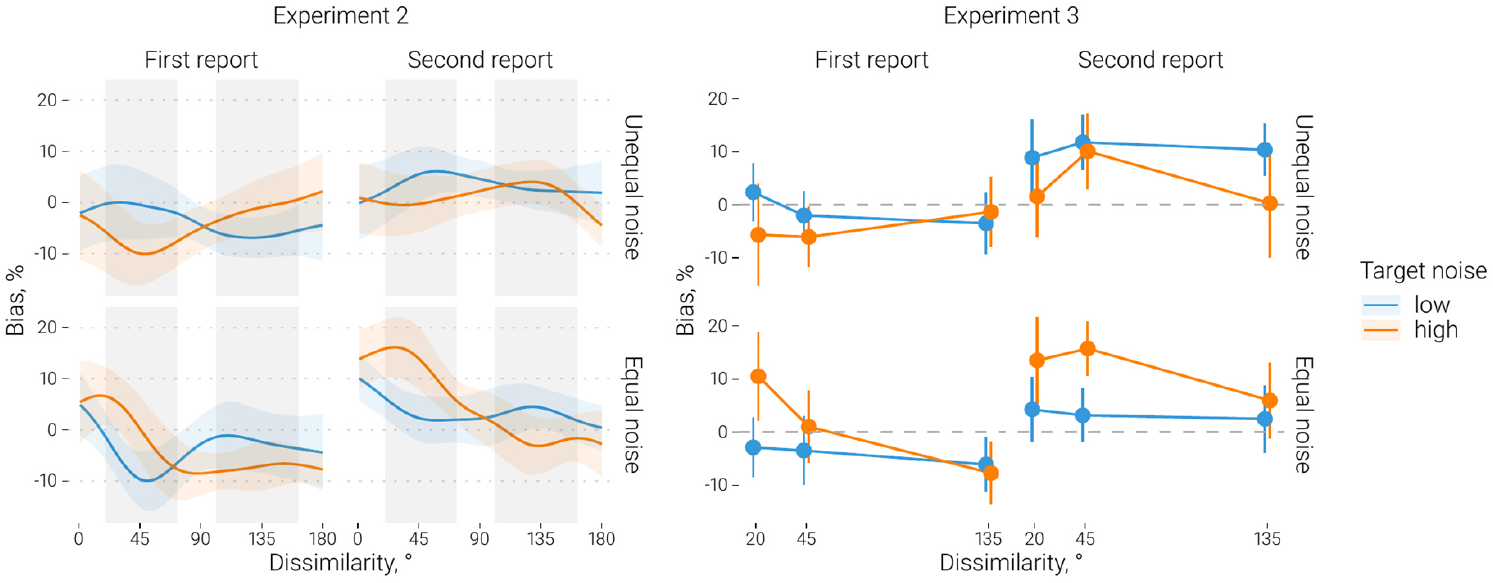
Biases within the first and the second response. In Experiments 2 and 3, we found that response order (first vs. second response) affects the overall balance between attraction and repulsion but does not interact with the other factors. In Experiment 2, the data from the first and the second response were used to create bias curves, which were then subject to the same analysis as the combined bias curves in the main text. The same two clusters with the interaction between target noise and noise equality as in the combined analyses were identified (from 20° to 73°, *p*_*perm*_ = .057; from 101° to 162°, *p*_*perm*_ = .031). However, no significant clusters that involved response order or its interactions with other factors were observed. Additional analyses indicated that, overall, biases in Experiment 2 were more attractive in the second response (*F*(1, 15) = 17.72, *p* < .001, *η*^2^_G_ = .17), but, as the figure shows, that overall shift did not change the effect of target noise or noise equality. In Experiment 3, we also repeated the analyses in the main text with the response order factor included. We found the same two interactions reported in the overall analysis, between target noise and noise equality (*F*(1, 22) = 29.19, *p* < .001, *η*^2^_G_ = .02) and between noise equality and the similarity bin (*F*(2, 36) = 3.66, *p* = .026, *η*^2^_G_ < .01). The effect of response order was also significant, with more attraction for the second response (*F*(1, 22) = 22.04, *p* < .001, *η*^2^_G_ = .06), but it did not interact with other factors. In sum, the second response shows more attraction, but the key interaction between the target noise and noise equality does not depend on it. Lines show the mean bias, and bars and shaded regions show 95% CIs.

**Figure S4.**
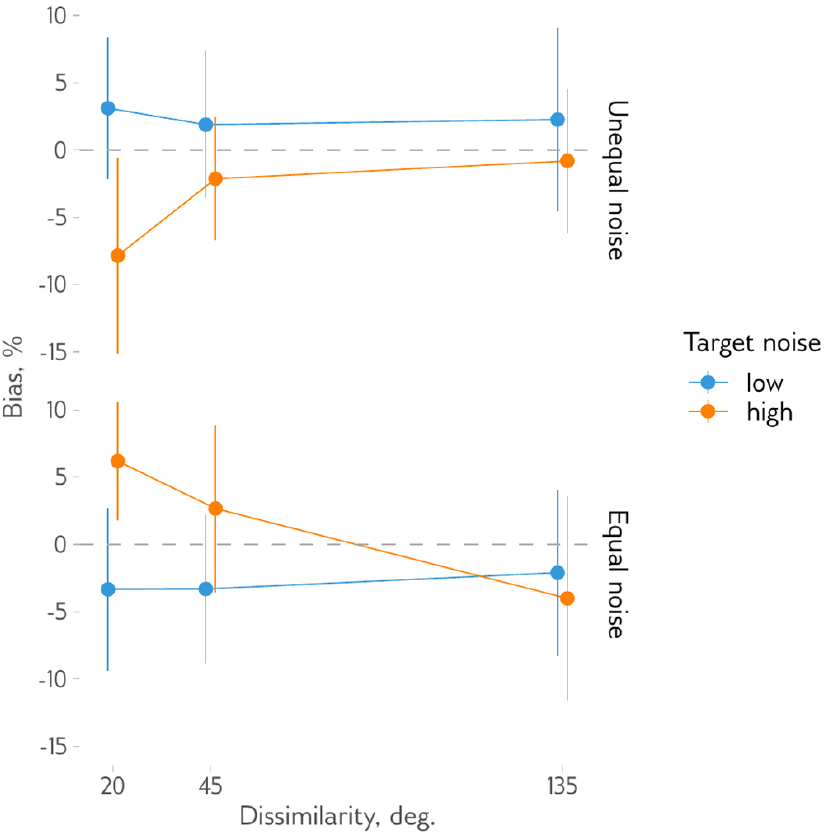
Biases in Experiment 3 for small (<10°) errors. As an additional test of the role of swap errors, we also repeated the analysis of biases but using only the errors below 10° in absolute value, which cannot be interpreted as swap errors even in the 20° similarity bin. Including only small errors in the analysis did not change the observed effects pattern, suggesting that the effect of target noise and noise equality on attractive biases in Exp. 3 is not likely to be explained by swap errors. Similar to the analyses of the full dataset, reported in the main text, we observed an interaction between the target noise level and noise equality (*F*(1, 22) = 7.82, *p* = .011, *η*^2^_G_ = .03), further qualified by a three-way interaction with the similarity bin (*F*(2, 44) = 3.80, *p* = .030, *η*^2^_G_ = .03).

